# Ancient retrotransposon-derived promoters for mammalian genomic imprinting

**DOI:** 10.1101/2025.07.14.664778

**Authors:** Joomyeong Kim

## Abstract

In mammalian genomic imprinting, RNA Pol II-driven transcription by upstream alternative promoters is responsible for establishing gametic DNA methylation on downstream Imprinting Control Regions (ICRs). According to surveys on germ cell transcripts, the alternative promoters for paternally expressed *PEG3*, *MEST* and *PLAGL1* are evolutionarily conserved and transcriptionally active mainly in the oocytes of the mammalian species. These promoters are usually bi-directional, producing transcripts in both directions. Furthermore, these alternative promoters are either part of or closely associated with the ancient retrotransposons that have inserted into the genomes more than 100 million years ago. The oocyte-specific alternative promoter of human *SNRPN* and the promoter of mammalian *AIRN* are also believed to have originated from the ancient transposon of similar evolutionary ages. These results are overall consistent with the well-known properties of retrotransposons, de-repression and transcriptional activity in germ cells. This further suggests that ancient retrotransposons may have been co-opted as *cis*-regulatory elements for establishing DNA methylation for genomic imprinting.

## Introduction

In mammalian genomes, a small subset of autosomal genes, about 200, are expressed mainly from one allele due to an epigenetic mechanism termed genomic imprinting, by which one allele is repressed through DNA methylation and histone modification [1, 2]. These imprinted genes are usually expressed in early-stage embryos, placenta and brain [1–3]. Mutations on these genes tend to cause changes in fetal growth rates and nurturing behaviors in the animals and various genetic disorders in humans [1, 2, 4]. Genomic imprinting is detected mainly in the eutherian lineage, which has an unusual reproduction scheme with viviparity and placentation. Thus, genomic imprinting may have co-evolved with this reproduction strategy of mammalian lineage [5–7].

Imprinted genes are usually clustered in specific regions of chromosomes, forming imprinted domains with the genomic size ranging from 0.5 to 2 mega base pair (Mb) in length. The imprinting or mono-allelic expression of each domain is regulated through small genomic regions termed Imprinting Control Regions (ICRs) [1, 2]. During gametogenesis, these ICRs acquire DNA methylation, which serves as a gametic signal for the next generation [1, 2]. According to the recent studies, DNA methylation on these ICRs is mediated through RNA Pol II-driven transcription involving alternative promoters that are located upstream of the ICRs. Alternative promoter-driven transcription is also accompanied with the histone modification H3K36me3 (Trimethylation on Lys36 of Histone 3) in the downstream regions, including the ICRs, which in turn recruits *de novo* DNA methylation machinery through the interaction of H3K36me3 with the PWWP (Pro-Trp-Trp-Pro) domain of DNMT3A (DNA methyltransferase 3A) [8–10]. This has been demonstrated through a series of experiments either deleting alternative promoters or truncating alternative transcription, which usually result in the loss of DNA methylation on a given ICR, subsequently disrupting the imprinting of the associated domain [11–17]. Thus, the alternative promoters are thought to be very critical for establishing gametic DNA methylation on ICRs [9, 14]. Nevertheless, several key features associated with these alternative promoters are currently unknown, such as their evolutionary origin and conservation among mammalian species.

To further characterize these alternative promoters, we previously performed a series of NGS (Next Generation Sequencing)-based 5’RACE (Rapid Amplification cDNA End) experiments, which have identified several alternative 1^st^ exons and promoters for individual imprinted genes in mouse, cow and human [18–20]. The current study has further extended this effort through surveying germ cell transcripts derived from the mammals. The results confirmed the evolutionary conservation of these alternative promoters in mammals. Interestingly, these promoters are all believed to have been derived from ancient retrotransposons. More details are described below.

## Materials and Methods

### Ethics statement

All the experiments related to mice were performed in accordance with National Institutes of Health guidelines for care and use of animals and also approved by the Louisiana State University Institutional Animal Care and Use Committee (IACUC), protocol #25-043.

### Bioinformatic approaches for the identification of alternative promoters

A large number of RNAseq data were initially retrieved from the GEO DataSets with the following key terms: RNAseq, mammals, oocytes or spermatocytes. This initial set of RNAseq data, 665 entries, were further filtered based on the depth and length of sequence reads per sample. The final set selected for this study is provided as **Supplementary file S1**. With the fasterq-dump command of sra-tools, individual SRR files were downloaded from the SRA Run Selector as fastq format files, which were then compressed into gz format files with the gzip command. A set of gz files for a given SRR file were used as input files with the mapping program HISAT2 against the indexed genome file that has been downloaded from NCBI [21]. An output sam format file was further sorted and converted into a bam format file with samtools. This process was executed with the following command line: hisat2 -q --rna-strandness R -x mm10 −1 SRRxxx_1.fastq.gz −2 SRRxxx_2.fastq.gz | samtools sort -o SRRxxx.bam. The index file for the final bam file was also produced with samtools with the following command line: samtools index SRRxxx.bam. The produced index file, SRRxxx.bam.bai, was stored in the same folder as the bam file to be uploaded onto the genome viewer IGV (Integrative Genomics Viewer) [22]. Finally, the exon structures of individual imprinted genes were manually inspected with IGV.

### Analyses of the repeat elements associated with alternative promoters

The genomic region surrounding the alternative promoter for a given imprinted gene was initially inspected through the Repeat Masker and Interrupted Repeat functions of the UCSC genome browser (https://genome.ucsc.edu/index.html). Subsequent detailed analyses were performed, individually, with the RepeatMasking program at the Repeat Masker Web Server (https://www.repeatmasker.org) [23]. The output from this individual analysis was further examined to confirm the joining or defragmentation of several repeat fragments located within a given genomic interval.

### Expression analyses through RT-PCR

A panel of cDNAs was generated through the total RNA isolated from the individual tissues of the adults and embryos of the mouse. The adult tissues were harvested from 2-month-old C57BL/6J males and females, and the 14.5-dpc (days post coitum) embryos and placentas were harvested from a pregnant female that has been timed-mated with an adult male mouse. The harvested tissues were used for isolating total RNA with the EasyBlue total isolation kit (Intron, Seoul, South Korea), which was then used for generating cDNA with the reverse transcription protocol utilizing the MMTV reverse transcriptase (New England Biolab). The quality of cDNA was further monitored through qRT-PCR measuring the amount of an internal control, β-actin.

Finally, the prepared set of cDNA was used to detect the expression of a given transcript. The detailed information regarding the sequence and position of each oligonucleotide primer is available from **Supplementary file S2**. The original images of RT-PCR products that were separated on 2% agarose gel electrophoresis are also provided as **Supplementary figure S1**.

## Results

### Alternative promoter of mammalian Peg3 (Paternally Expressed Gene 3)

In the current study, we decided to re-analyze available transcriptome data that were derived from the germ cells of several mammalian species, including human and crab-eating macaque for the primates, mouse and rat for the rodents, cow and pig for the artiodactyls. We have downloaded this set of RNAseq data from the NCBI GEO database, which were generated from the total RNA isolated from Metaphase II (M2) oocytes and spermatocytes or purified prospermatogonia (**Supplementary file S1**). These data sets were subsequently processed through bioinformatic pipelines to map individual transcripts to genomes, the mapped transcriptomes were uploaded onto the visualization tool IGV (Integrative Genomics Viewer), and finally the exon structures of individual imprinted genes were manually inspected to identify potential alternative promoters.

First, the mouse *Peg3* locus was examined with the mapped transcripts derived from M2 oocytes (**Figure 1A**) and from prospermatogonia (**Supplementary figure S2**). According to the previous studies, the imprinting of mouse *Peg3* is controlled through the Peg3-DMR (Differentially Methylated Region) as an ICR, which encompasses the 4-kb genomic region containing the first exons of both *Peg3* and *Usp29* (Ubiquitin-Specific Protease 29) (E1 in **Figure 1A**) [24–26]. This imprinted locus is known to contain three alternative promoters/exons, U1, U2 and U3, that are located 19, 26, and 160-kb upstream of the E1 promoter. Yet, U1 is the only one that is highly expressed in oocytes [18, 20]. Consistent with this, the mapped transcripts of *Peg3* from M2 oocytes all start from the alternative promoter/exon U1, but not from the main promoter/exon E1, which is transcriptionally active in somatic cells. These transcripts are also connected to the downstream alternative exon U0 as shown in the splice junction of the transcriptome. This series of analyses also revealed that this alternative U1 promoter is bi-directional, triggering transcription in the opposite direction. This transcript is named as Oocyte-Specific Transcript (*Ost*), which is made of two exons that are separated by a 15-kb genomic distance. According to the results from prospermatogonia, the E1 promoter is the main one triggering transcription in both directions, producing the transcripts for both *Peg3* and *Usp29*. Interestingly, the alternative promoter U1 also triggers transcription in both directions, but with a much weaker strength than the main promoter E1 based on the number of the mapped transcripts (**Supplementary figure S2**).

**Figure 1.**
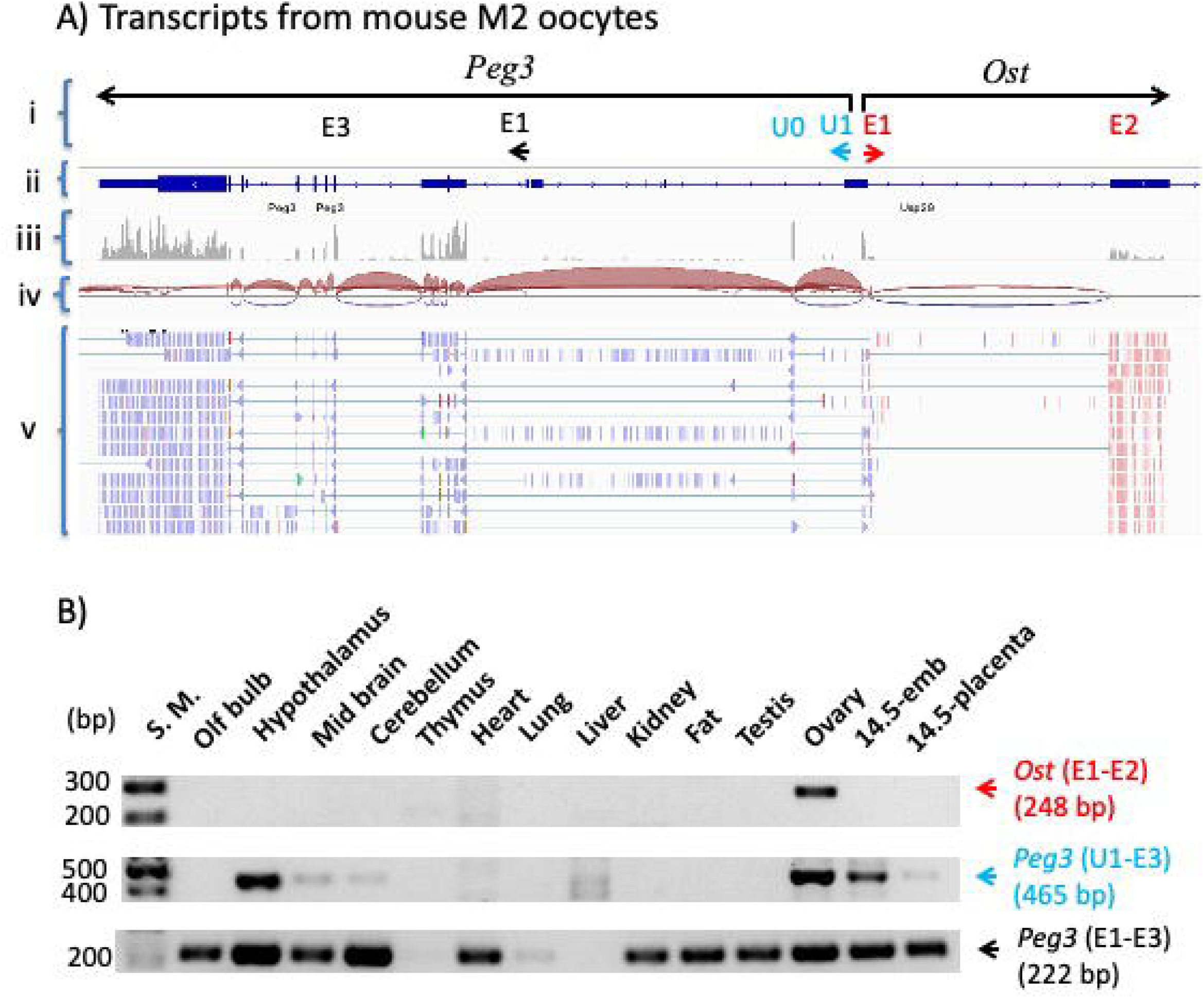
Alternative promoter/exon (U1) of the mouse *Peg3* locus. **(A)** Mapped transcripts derived from M2 oocytes (SRR2072743/GSE70116) were visualized with IGV (Integrative Genomics Viewer): the first row (i) representing the custom annotation based on the current study; the second row (ii) representing the NCBI annotation derived from the available public data; the third row (iii) representing the genomic coverage by the mapped transcripts from RNAseq; the fourth row (iv) representing the splice-junction derived from the mapped transcripts; the fifth row (v) representing the alignment of individual sequence reads against the genome sequence. Individual sequence reads are also represented with either red or blue lines to indicate the two different directions of the transcripts. In the first row (i), the transcriptional direction of *Peg3* is indicated with an arrow, and the positions of an alternative promoter/exon U1 and downstream another exon U0 are shown underneath the arrow. The positions of the somatic promoter/first exon (E1) and the third exon (E3) are also indicated. The transcript *Ost* in the opposite direction by the alternative promoter U1 is indicated with another arrow and its two exons (E1 and E2). **(B)** RT-PCR-based expression analyses. A panel of 14 cDNAs prepared from the individual tissues of 2-month-old adult mouse and 14.5-dpc embryo and placenta was used to survey the expression pattern of each transcript. Each set of primers and the size of the corresponding PCR product are indicated within two parentheses next to the name of the transcript. The relative position of each primer is also shown as part of the exon structure described above.

The expression pattern of the identified U1 promoter was further analyzed with a panel of cDNAs that have been prepared from the mouse tissues of adults and embryos (**Figure 1B**). As expected, the expression of *Peg3* was detected at high levels in the majority of the tissues, which is driven by the somatic promoter E1 (Product from E1-E3). The transcript starting from U1 was detected mainly in the ovary and hypothalamus of adults, and also at some levels in the 14.5-dpc (days post coitum) embryo (Product U1-E3). The high-level detection in the ovary is consistent with the fact that a large number of transcripts from M2 oocytes start from the U1 promoter, but not from the E1 promoter. This is also the case for Peg3-Ost, displaying its expression only in the ovary (Product E1-E2). Overall, this series of expression analyses confirmed the bi-directional promoter activity of U1 in the ovary.

The evolutionary conservation of the alternative promoter U1 was further tested through analyzing the transcriptome data derived from the germ cells of the other mammals, including rat, human, macaque, cow and pig. The data from the other mammals showed overall similar outcomes as the mouse, but with some species-specific differences. The data sets from human and cow are presented (**Figure 2** and **Supplementary figure S5**). As shown in **Figure 2A**, the human *PEG3* locus also contains the U1 promoter, which triggers transcription in both directions. Interestingly, the strength of the human U1 promoter appears to be much weaker than that observed from the mouse, which is based on the relative numbers of mapped transcripts.

**Figure 2.**
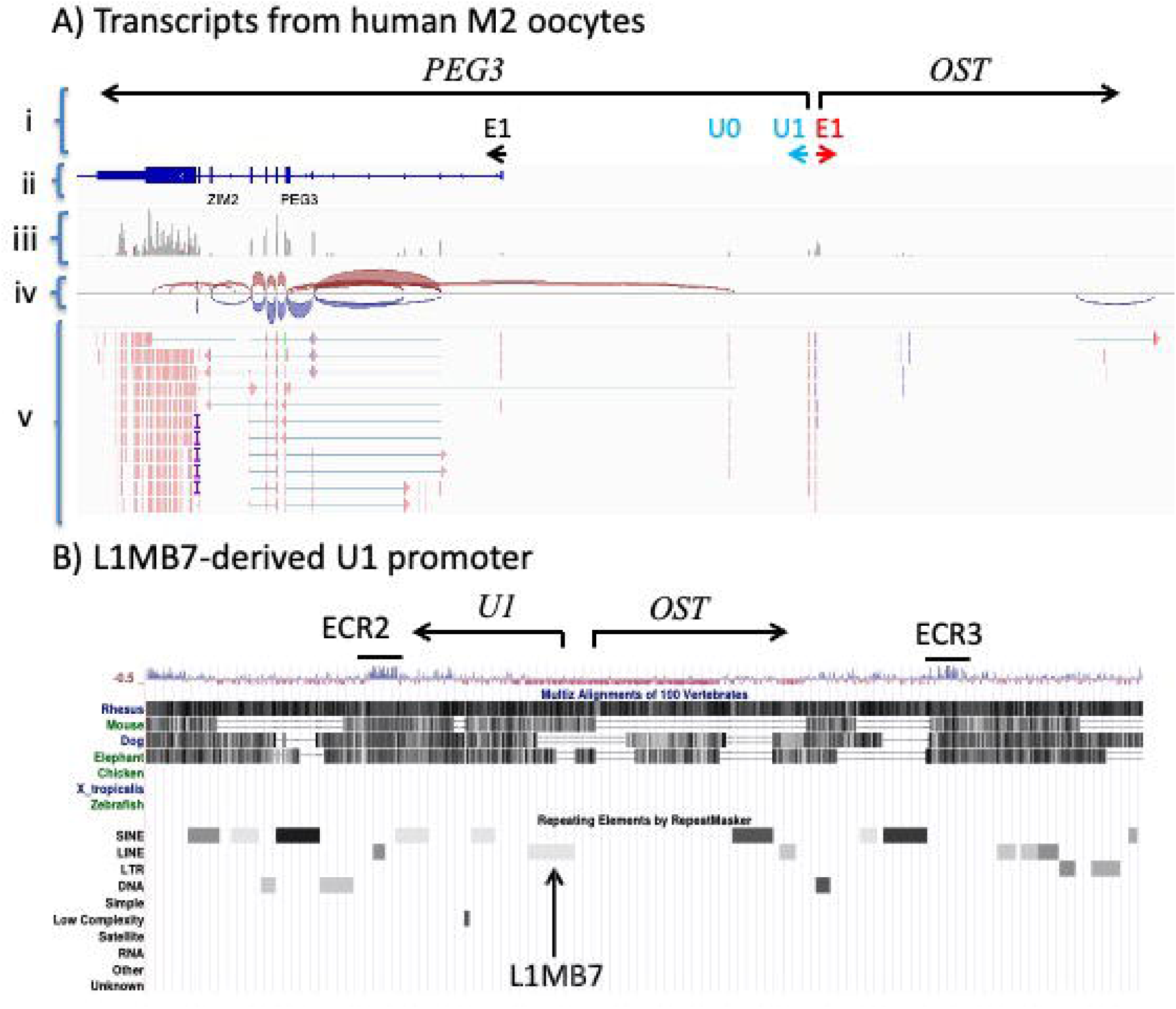
Alternative promoter/exon (U1) of the human *PEG3* locus. **(A)** Mapped transcripts derived from M2 oocytes (SRR12642767/GSE157834) were visualized with IGV (Integrative Genomics Viewer): the first row (i) representing the custom annotation based on the current study; the second row (ii) representing the NCBI annotation derived from the available public data; the third row (iii) representing the genomic coverage by the mapped transcripts from RNAseq; the fourth row (iv) representing the splice-junction derived from the mapped transcripts; the fifth row (v) representing the alignment of individual sequence reads against the genome sequence. Individual sequence reads are also represented with either red or blue lines to indicate the two different directions of the transcripts. In the first row (i), the transcriptional direction of *PEG3* is indicated with an arrow, and the positions of an alternative promoter/exon U1 and downstream another exon U0 are shown underneath the arrow. The position of the somatic promoter/first exon (E1) is also shown underneath the arrow. The transcript *OST* in the opposite direction by the alternative promoter U1 is indicated with an arrow. **(B)** Repeat element profile of the genomic region surrounding the human U1 promoter. The transcriptional start sites and directions of both U1-driven *PEG3* and *OST* transcripts are indicated with two arrows. The sequence conservation profile is also shown along with the repeat element profile. The U1 promoter region overlaps with the genomic region occupied by an ancient repeat element (L1MB7), which is indicated with a vertical arrow. This U1 promoter is also localized between two ECRs (Evolutionarily Conserved Regions), ECR2 and ECR3.

Also, the exon-joining of the U1 transcript is slightly different from that of the mouse: the transcript from U1 or U0 is spliced to the third exon (E3), but not to the second exon (E2) of *PEG3*. The strength of the U1 promoter in the direction of *Ost* was also weaker than that of the mouse given a small number of mapped transcripts. Similarly, the lengths and exon numbers of the *Ost* transcripts are variable among individual mammals: the longest one with 3 exons covering a 63-kb genomic distance in cow versus the shortest one with two exons covering a 15-kb genomic distance in mouse (**Figure 1A** and **Supplementary figure S5**). Overall, this series of analyses confirmed the evolutionary conservation of the bi-directional promoter U1 among mammalian species, but with some species-specific differences.

The sequence of the genomic region surrounding the mammalian U1 promoter was also carefully examined. As shown in **Figure 2B**, the human U1 promoter is flanked by two patches of evolutionarily conserved regions, ECR2 and ECR3, which are putative transcriptional enhancers based on their association with the histone modification H3K4me1 [27]. This genomic region is also occupied by a set of ancient repeats that have inserted into the genome. The sequence divergence of these ancient elements is represented through the varying degree of gray color in the Repeat element profiles (the lighter gray corresponding to the higher divergence with the older evolutionary age in **Figure 2B**). Among these repeat elements, an ancient L1 element belonging to the L1MB7 family is colocalized with the human U1 promoter. This colocalization is also detected in the other mammals, including macaque, rabbit, dog, elephant, and cow (**Supplementary figure S5**). However, this colocalization was not detected in some of the rodents, mouse and rat, which might be caused by the faster neutral mutation rates of the rodents than the other mammals, causing the rapid decay of retrotransposons in the rodent genomes [28]. Overall, the genomic layout surrounding the U1 promoter is well conserved among mammals, in particular the colocalization of U1 with the ancient L1, further suggesting that this ancient L1 might have been selected as part of the alternative promoter U1 for the mammalian *Peg3* locus.

### Alternative promoters of mammalian Mest (Mesoderm-Specific Transcript)

The mouse *Mest* locus has been analyzed similarly as the *Peg3* locus (**Figures 3** and **4**). According to the previous study, the mouse *Mest* locus is known to have two alternative promoters/exons, U1 and U2, which are located 5-kb and 15-kb upstream of the main somatic promoter/exon E1 [20]. This series of transcriptome analyses further identified an additional alternative promoter/exon, U3, which is located 30-kb upstream of the E1 promoter (**Figure 3A**). This alternative promoter also triggers transcription in both directions, thus the alternative exon in either direction is joined to the 2^nd^ exon of the two flanking genes, *Mest* and *Cep41* (Centrosomal protein 41). The transcription from this bi-directional promoter were further tested through a series of expression analyses employing the same panel of mouse cDNA as the *Peg3* locus (**Figure 3B**). As expected, the high-level expression of mouse *Mest* was detected in the various somatic tissues (E1-E2 in **Fig 3B**). In contrast, the transcripts in both directions by the U3 promoter were detected mainly in the ovary and at some levels in the 14.5-dpc embryo (U3-E2 for the *Mest* direction and U1-E2 for the *Cep41* direction). The promoter activity of U3 was not detected in the prospermatogonia with the majority of the mapped transcripts starting from the E1 promoters for *Mest* and *Cep41* (**Supplementary figure S3**). Thus, this series of analyses confirmed the bi-directional promoter activity of U3 mainly in ovary.

**Figure 3.**
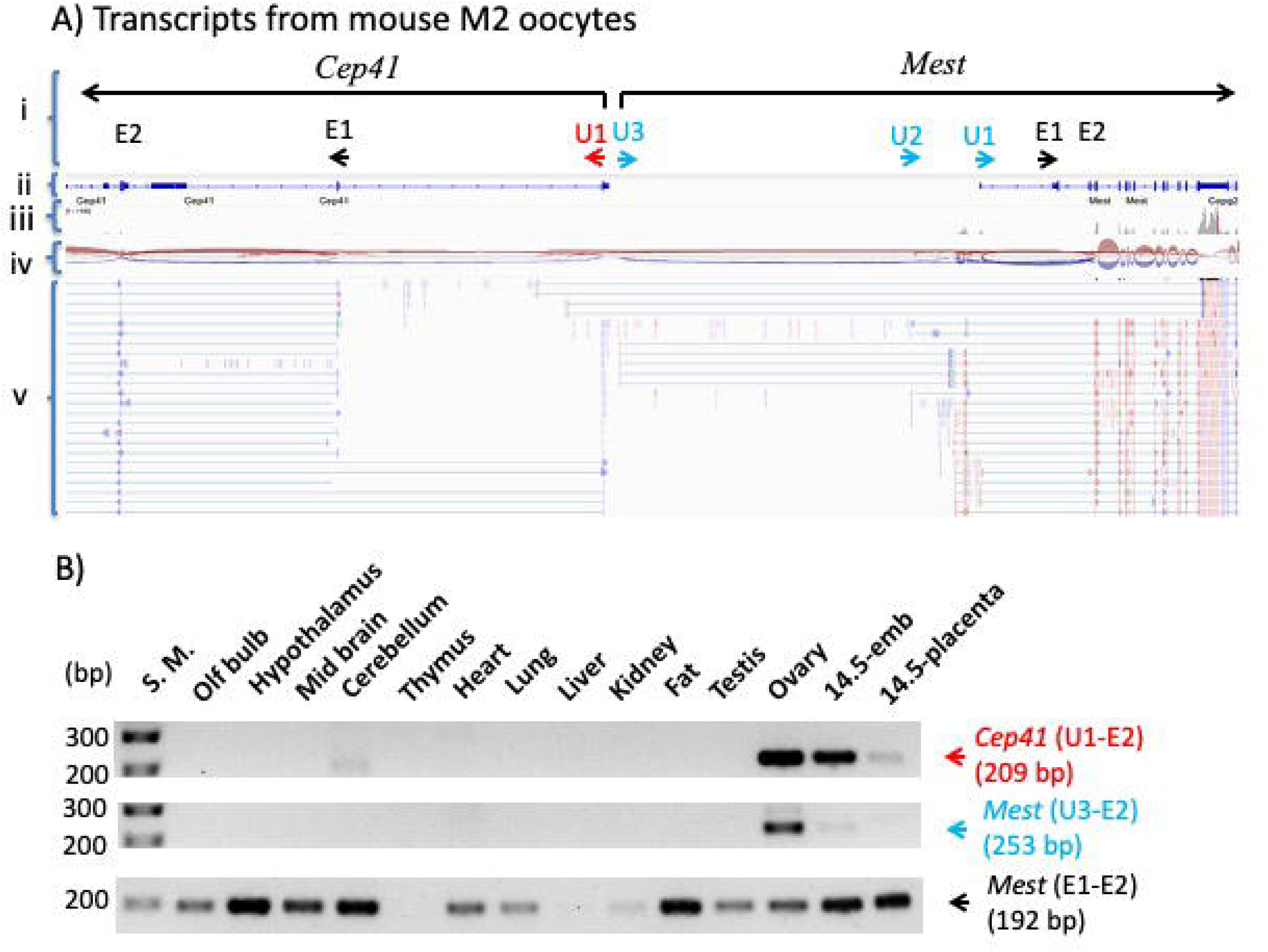
Alternative promoters/exons (U3-U1) of the mouse *Mest* locus. **(A)** Mapped transcripts derived from M2 oocytes (SRR2072743/GSE70116) were visualized with IGV (Integrative Genomics Viewer): the first row (i) representing the custom annotation based on the current study; the second row (ii) representing the NCBI annotation derived from the available public data; the third row (iii) representing the genomic coverage by the mapped transcripts from RNAseq; the fourth row (iv) representing the splice-junction derived from the mapped transcripts; the fifth row (v) representing the alignment of individual sequence reads against the genome sequence. Individual sequence reads are also represented with either red or blue lines to indicate the two different directions of the transcripts. In the first row (i), the transcriptional direction of *Mest* is indicated with an arrow, and the positions of alternative promoters/exons (U3-U1) are also shown underneath the arrow. The positions of the somatic promoter/first exon (E1) and the second exon (E2) are indicated. The transcription in the opposite direction also starts from the alternative promoter U3 towards the adjacent gene *Cep41*, which is indicated with another arrow along with the alternative promoter/exon U1 and two downstream exons (E1 and E2). **(B)** RT-PCR-based expression analyses. A panel of 14 cDNAs from the individual tissues of 2-month-old adult mouse and 14.5-dpc embryo and placenta was used to survey the expression pattern of each transcript. Each set of primers and the size of the corresponding PCR product are indicated within two parentheses next to the name of the transcript. The relative position of each primer is also presented as part of the exon structure described above.

**Figure 4.**
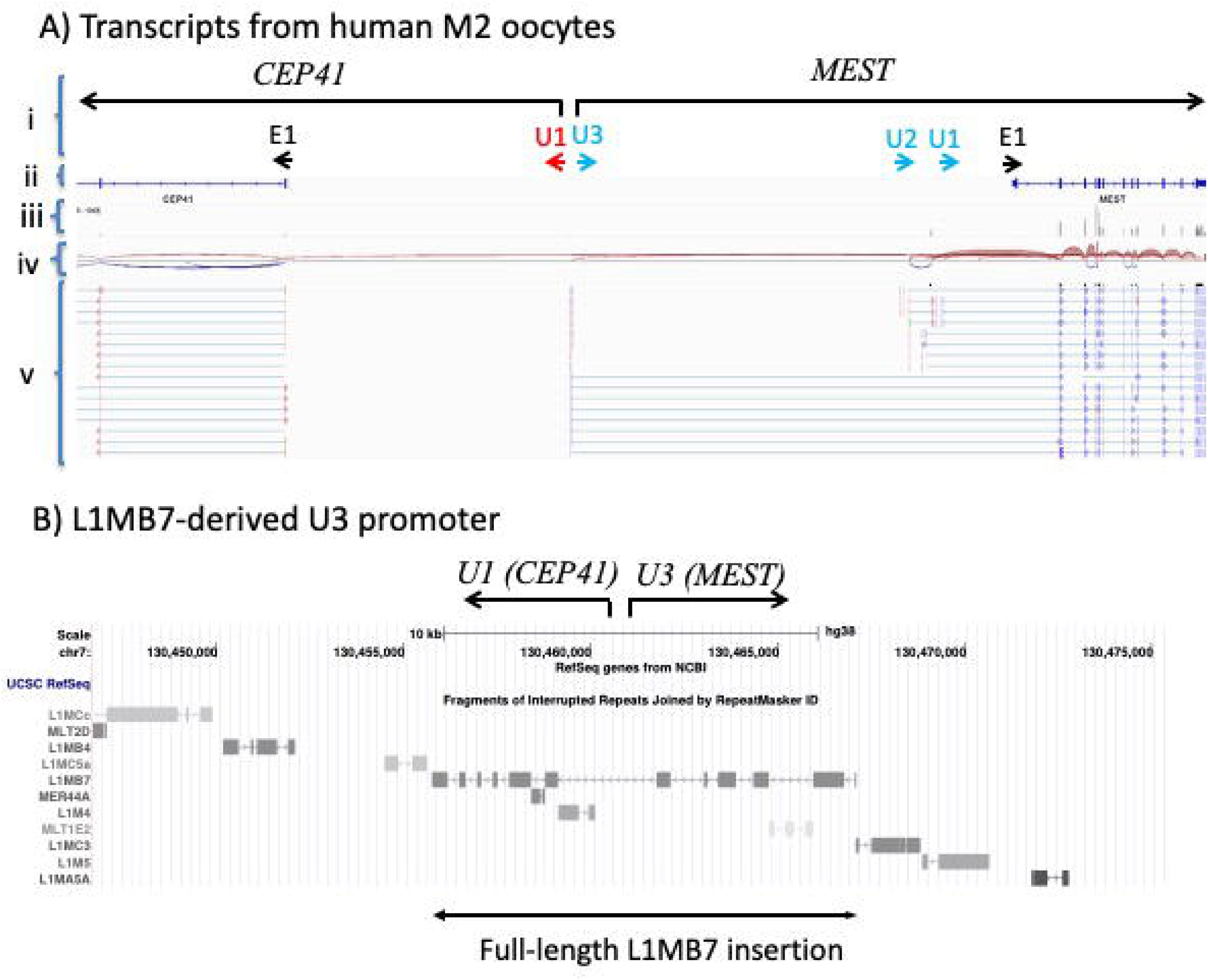
Alternative promoters/exons (U3-U1) of the human *MEST* locus. **(A)** Mapped transcripts derived from M2 oocytes (SRR12642767/GSE157834) were visualized with IGV (Integrative Genomics Viewer): the first row (i) representing the custom annotation based on the current study; the second row (ii) representing the NCBI annotation derived from the available public data; the third row (iii) representing the genomic coverage by the mapped transcripts from RNAseq; the fourth row (iv) representing the splice-junction derived from the mapped transcripts; the fifth row (v) representing the alignment of individual sequence reads against the genome sequence. Individual sequence reads are also represented with either red or blue lines to indicate the two different directions of the transcripts. In the first row (i), the transcriptional direction of *MEST* is indicated with an arrow and the positions of alternative promoter/exon (U3-U1) are also shown underneath the arrow. The position of the somatic promoter/first exon (E1) is indicated with a small arrow. The transcription in the opposite direction towards the adjacent gene *CEP41* is indicated by an arrow along with an alternative promoter/exon U1. **(B)** Repeat element profile of the genomic region surrounding the human U3 promoter. The transcriptional start sites and directions of both U3-driven *MEST* and U1-driven *CEP41* transcripts are indicated by two arrows. The 10-kb genomic region surrounding the transcription sites is represented by a set of 11 ancient repeat elements, which are believed to have been derived from the insertion and subsequent fragmentation of a full-length L1 element, L1MB7.

The evolutionary conservation of the alternative promoter U3 was further tested through analyzing the transcriptomes derived from the other mammals, including human (**Figure 4A**) and cow (**Supplementary figure S6**). As seen in the mouse, the mammalian U3 promoter was also found to be localized along with the two other alternative promoters, U1 and U2, with their similar relative genomic positions to the somatic promoter E1. The mammalian U3 promoter also triggers transcription in both directions. However, none of the transcripts in the direction of *Cep41* are joined to the second exon of *Cep41*, which is different from the exon-joining detected from the mouse. The genomic sequence surrounding the mammalian U3 promoter were also analyzed in terms of their repeat element profiles (**Figure 4B** and **Supplementary figure S6**).

According to the results, the 10-kb genomic region surrounding the U3 promoter contains a series of repeat elements that are recognized as part of the ancient retrotransposon family L1MB7, which is the same L1 family as detected from the U1 promoter of mammalian *Peg3* (**Figure 2B**). Furthermore, the entire set of twelve L1MB7-derived fragments could be joined back together as one full-length L1 element based on the position and orientation of each fragment relative to an intact full-length L1 element [29]. Thus, it is believed that the insertion and subsequent fragmentation might have initiated or provided a molecular environment for the formation of the current U3 promoter for the mammalian *Mest* locus.

### Alternative promoter of mammalian Plagl1 (PLAG1 Like zinc finger 1)

According to the previous studies, the mouse *Plagl1* locus is also known to have an alternative promoter/exon, U1, which is located 30-kb upstream of the somatic promoter/exon E1 (**Figure 5A**) [20]. This series of analyses indeed confirmed the promoter activity of U1 in M2 oocytes with the majority of transcripts starting from this alternative promoter. Also, this U1 promoter appears to be bi-directional, triggering another transcript, *Ost*, in the opposite direction, which is similar to the other loci described above (**Figure 5A**). Subsequent expression analyses confirmed the expression of both transcripts in ovary as well as in thymus (U1-E3 for *Plagl1* and E1-E2 for *Ost* in **Figure 5B**). Some levels of the promoter activity were also detected in the prospermatogonia, but with much weaker strength, and also none of the transcripts was spliced to the exons of *Plagl1* (**Supplementary figure S4**).

**Figure 5.**
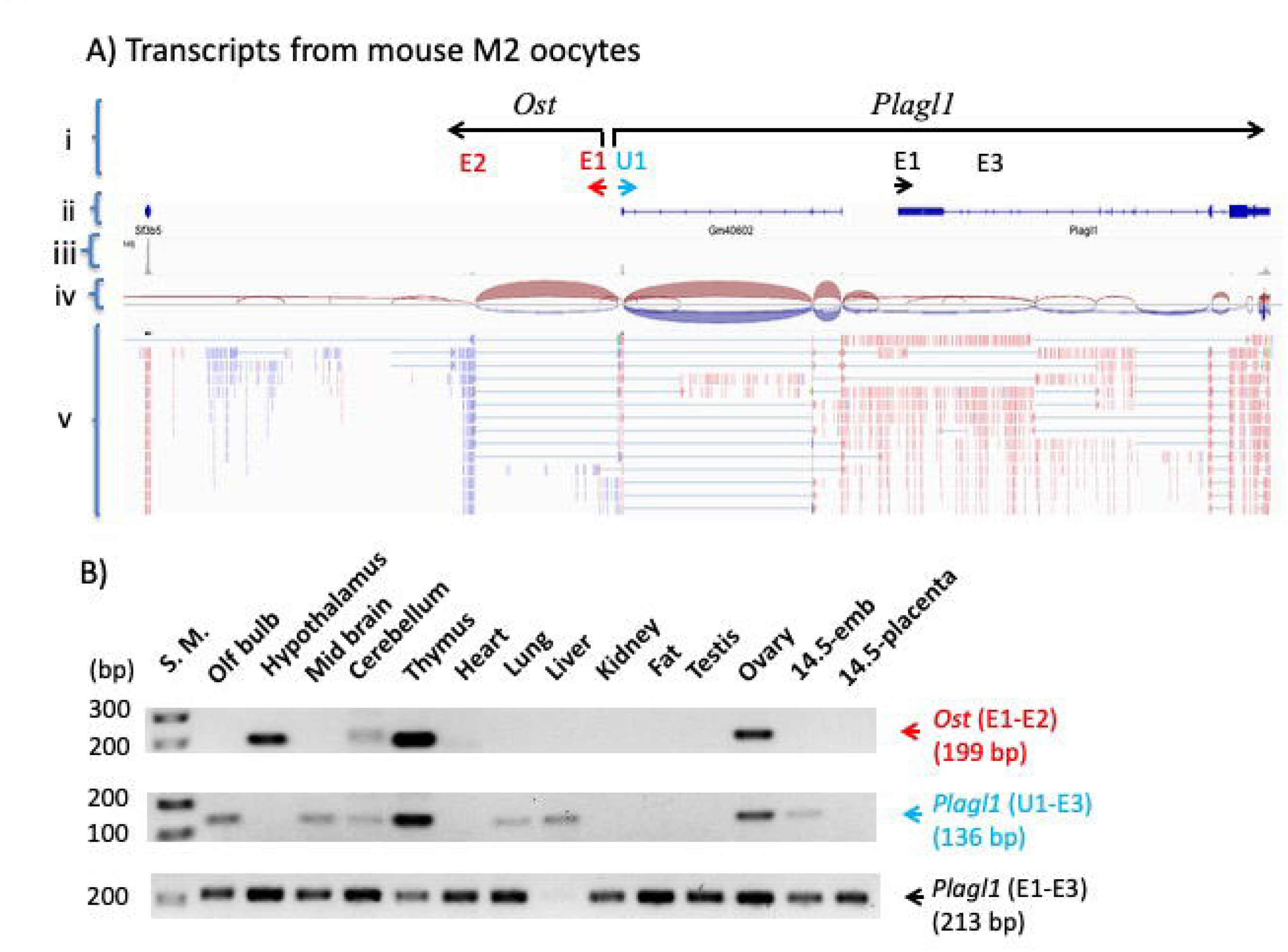
Alternative promoter/exon (U1) of the mouse *Plagl1* locus. **(A)** Mapped transcripts derived from M2 oocytes (SRR2072743/GSE70116) were visualized with IGV (Integrative Genomics Viewer): the first row (i) representing the custom annotation based on the current study; the second row (ii) representing the NCBI annotation derived from the available public data; the third row (iii) representing the genomic coverage by the mapped transcripts from RNAseq; the fourth row (iv) representing the splice-junction derived from the mapped transcripts; the fifth row (v) representing the alignment of individual sequence reads against the genome sequence. Individual sequence reads are also represented with either red or blue lines to indicate the two different directions of the transcripts. In the first row (i), the transcriptional direction of *Plagl1* is indicated with an arrow, and the positions of the alternative promoter/exon U1, the somatic promoter/first exon (E1) and the third exon (E3) are also shown. The transcript *Ost* in the opposite direction by the alternative promoter U1 is indicated with an arrow, which is made of two exons (E1 and E2). **B)** RT-PCR-based expression analyses. A panel of 14 cDNAs from the individual tissues of 2-month-old adult mouse and 14.5-dpc embryo and placenta was used to survey the expression pattern of each transcript. Each set of primers and the size of the corresponding PCR product are indicated within two parentheses next to the name of the transcript. The relative position of each primer is also shown as part of the exon structure described above.

The evolutionary conservation of the U1 promoter was also tested through analyzing the transcriptomes of the other mammals. According to the results, the other mammals indeed have this alternative promoter as the main promoter triggering the transcription for mammalian *Plagl1* in M2 oocytes (human in **Figure 6A** and cow in **Supplementary figure S7**). However, the transcript in the opposite direction, *Ost*, was not detected in the M2 oocytes of human, while detected in cow. Detailed examination further revealed that the transcription starts at the two different locations for human *PLAGL1* (U1.1 and U1.2 in **Figure 6B**). The genomic region surrounding one of these start sites (U1.1) appears to be located within a 160-bp genomic distance to a repeat element belonging to the retrotransposon family MIR (Mammalian-wide Interspersed Repeat), which is one of ancient retrotransposon families [30–32]. This MIR element is conserved among all the mammals and, in fact, recognized as part of a proximal enhancer by the ENOCODE project [33]. In cow, this MIR element is again the main alternative promoter for the direction of *Plagl1*, yet another adjacent MIR element triggers transcription in the opposite direction, producing *Ost* (**Supplementary figure S7**). Overall, all the mammals appear to contain the alternative promoter U1 for the *Plagl1* locus, which is closely associated with another ancient retrotransposon element belonging to the MIR family. This close association further suggests potential involvement of retrotransposons in the formation of the alternative promoter for the mammalian *Plagl1* locus.

**Figure 6.**
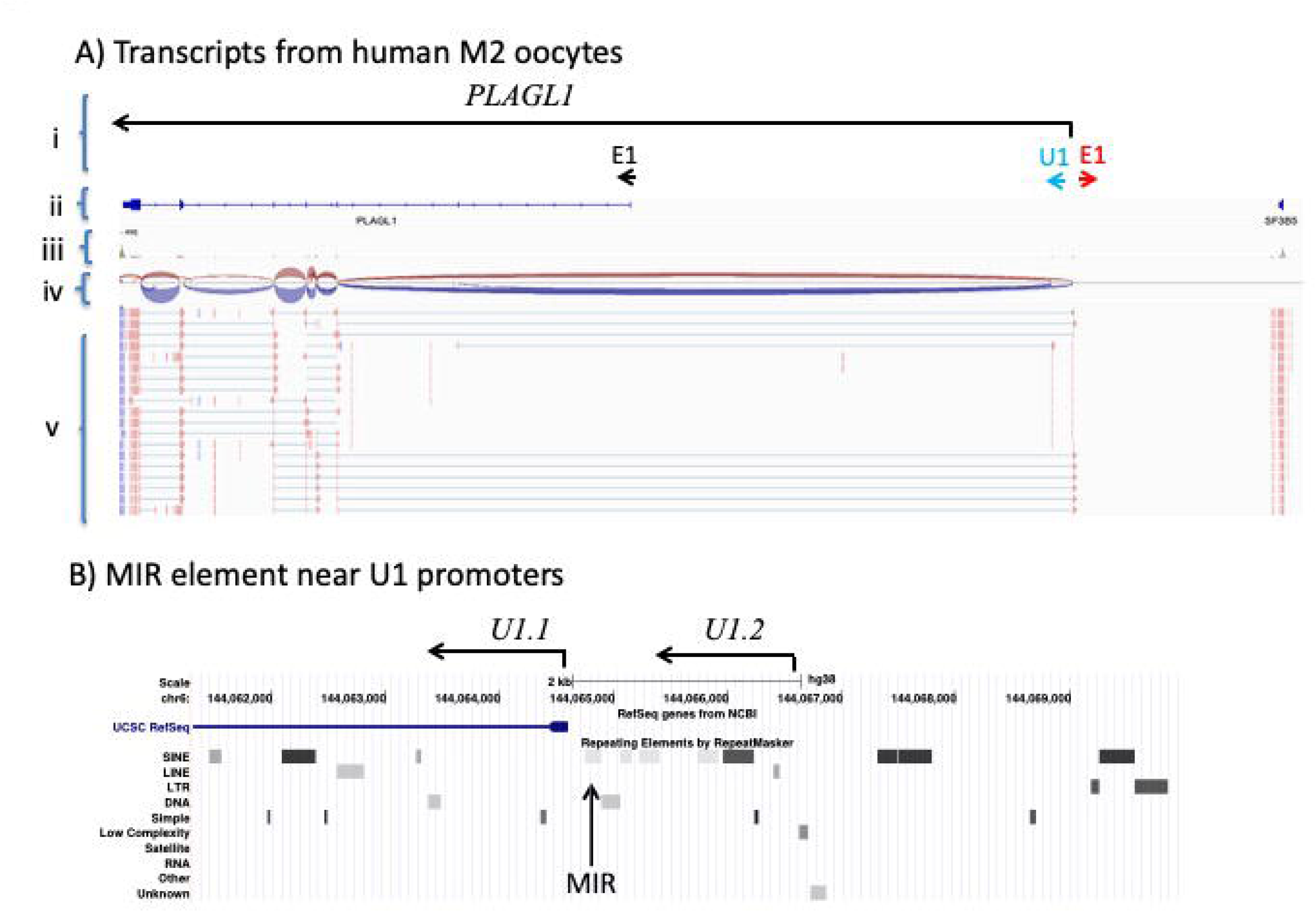
Alternative promoter/exon (U1) of the human *PLAGL1* locus. **(A)** Mapped transcripts derived from M2 oocytes (SRR12642767/GSE157834) were visualized with IGV (Integrative Genomics Viewer): the first row (i) representing the custom annotation based on the current study; the second row (ii) representing the NCBI annotation derived from the available public data; the third row (iii) representing the genomic coverage by the mapped transcripts from RNAseq; the fourth row (iv) representing the splice-junction derived from the mapped transcripts; the fifth row (v) representing the alignment of individual sequence reads against the genome sequence. Individual sequence reads are also represented with either red or blue lines to indicate the two different directions of the transcripts. In the first row (i), the transcriptional direction of *PLAGL1* is indicated with an arrow, and the positions of the alternative promoter/exon U1 and the somatic promoter/first exon (E1) are also shown underneath the arrow. The transcript *OST* in the opposite direction by the alternative promoter U1 is not detected, despite a small number of mapped transcripts in the opposite direction (E1). **(B)** Repeat element profile of the genomic region surrounding the human U1 promoters. The transcriptional direction is indicated with the two arrows in the same direction to represent the two different start sites of the U1-driven *PLAGL1* transcripts (U1.1 and U1.2). One of these two start sites is located very close to an MIR element as indicated by a vertical arrow.

### Ancient retrotransposons associated with mammalian Snrpn and Airn

Given the observations described above, we decided to perform another series of thorough inspection to test whether the promoters of additional imprinted loci are also closely associated with ancient retrotransposons. According to this survey, the promoters of two imprinted loci were found to be colocalized with ancient retrotransposons (**Figure 7**). First, the human *SNRPN* (Small nuclear ribonucleoprotein polypeptide N) is known to contain a set of 11 alternative promoters that are localized within the 130-kb genomic region upstream of its somatic main promoter. Among these upstream alternative promoters, two promoters are oocyte-specific (U5 and U6) and responsible for DNA methylation setting on the main promoter of *SNRPN* during oogenesis [13]. In fact, the 880-bp region encompassing these two promoters was initially identified as AS-SRO (Angelman Syndrome Smallest Region of Deletion Overlap) (**Figure 7A**). In this case of Angelman syndrome, the deletion of this 880-bp region is thought to be responsible for loss of DNA methylation on the main promoter of *SNRPN* with biallelic expression, which in turn causes loss of the maternal expression of *UBE3A* (Ubiquitin protein ligase E3A) [13]. Yet, the U5 promoter turns out to be colocalized with a member of the ancient DNA transposon family MER5A (Medium Reiterated Sequence 5A) (**Figure 7A**). This ancient repeat is also detected in the upstream region of the other mammalian *Snrpn* locus, including squirrel, rabbit, dog, elephant and cow (**Supplementary figure S8**). Thus, this suggests that this ancient transposon might have been selected as an alternative promoter for mammalian *Snrpn*.

**Figure 7.**
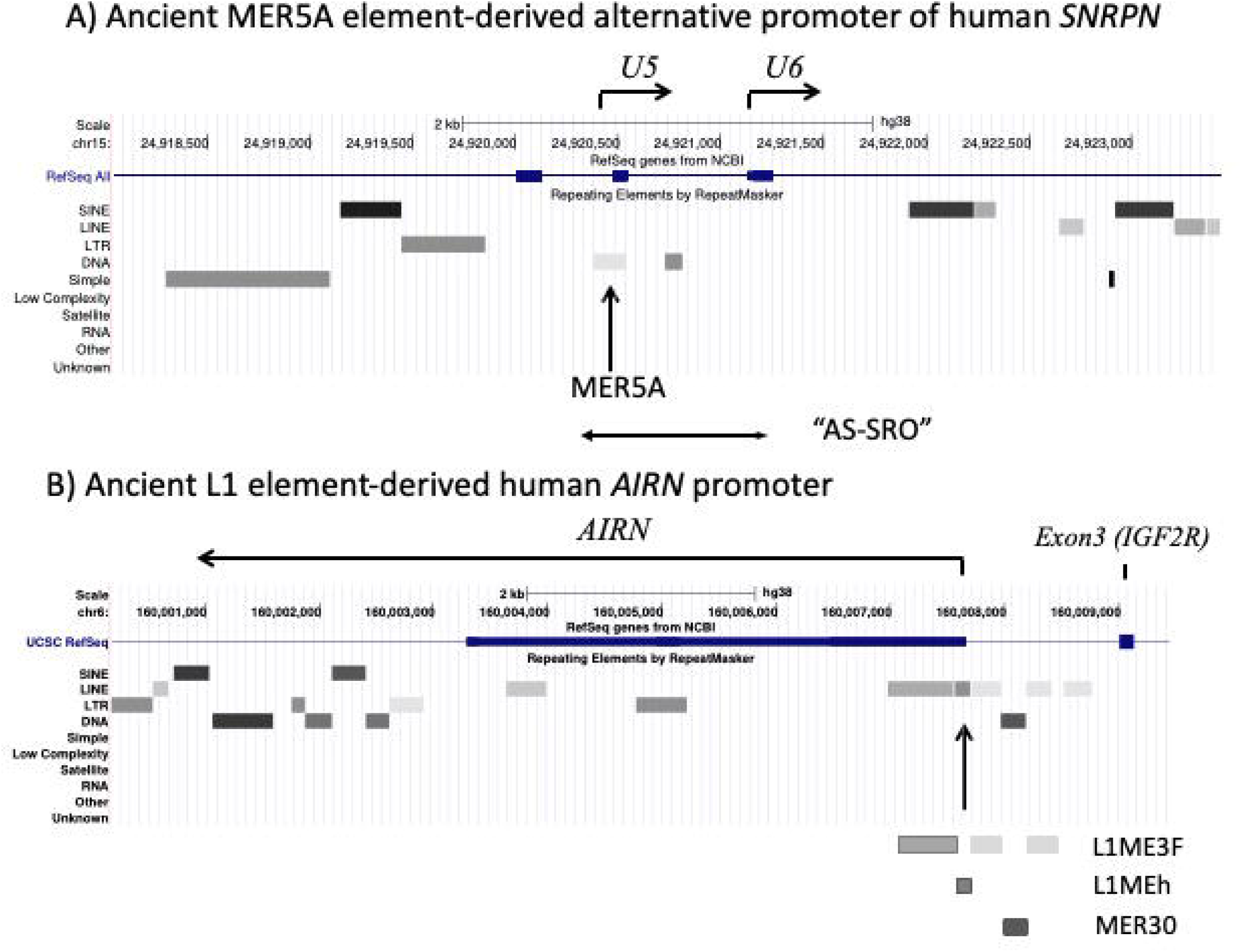
Ancient repeat elements associated with the promoters of human *SNRPN* and *AIRN*. **(A)** The genomic region surrounding the two oocyte-specific alternative promoters (U5 and U6) of human *SNRPN* contains several repeat elements. The U5 promoter is colocalized with a repeat element belonging to the transposon family MER5A. The 880-bp region encompassing U5 and U6 was previously identified as AS-SRO (Angelman Syndrome Smallest Region of Deletion Overlap), which is indicated with a horizontal line. **(B)** The transcription of human *AIRN* starts from the genomic region within the 2^nd^ intron of *IGF2R* in an opposite direction, which iis indicated with an arrow. The genomic region surrounding the promoter of human *AIRN* is filled with several repeat elements. These repeat elements are believed to have been derived from the initial insertion of L1ME3F, and later individually by L1MEh and DNA transposon MER30. The transcription of *AIRN* starts within L1MEh, which is also found in the other mammals.

Second, the mammalian *Igf2r* (Insulin like growth factor 2 receptor) locus contains another evolutionarily conserved gene, *Airn* (Antisense of Igf2r Non-coding RNA), yet the transcription of this antisense RNA gene is known to start from the 2^nd^ intron of *Igf2r* in the opposite direction (**Figure 7B**). The genomic region surrounding this promoter is also known to be a DMR (Differentially Methylated Region), which acquires gametic DNA methylation during oogenesis [34]. According to careful inspection, this 1.8-kb genomic region of the human *AIRN* promoter is filled with several repeat elements that belong to the ancient retrotransposon family L1ME3F. These elements could be joined back together into one long element based on their relative positions and orientations compared to its intact element [29]. Two subsequent insertions are thought to have occurred, individually, fragmenting the initial element of L1ME3F into three pieces, one insertion by a repeat element belonging to the L1MEh family and the other by an element belonging to the DNA transposon family MER30 (Medium Reiterated Sequence 30) (**Figure 7B**). The transcription of human *AIRN* is known to start within the inserted element of L1MEh, yet this element is detected among all the other mammals (**Supplementary figure S8**), suggesting that this element might have been selected as the promoter for this antisense Non-coding RNA gene to the mammalian *Igf2r* locus. Overall, the observed frequent associations suggest that retrotransposons most likely play significant roles in the formation and evolution of mammalian imprinted loci.

## Discussion

The current study reports that the alternative promoters for paternally expressed *PEG3*, *MEST* and *PLAGL1* are evolutionarily conserved and transcriptionally active mainly in the oocytes of all the mammalian species. These promoters are bi-directional, producing transcripts in both directions (**Figures 1-6**). Furthermore, these alternative promoters are either part of or closely associated with the ancient retrotransposons that have inserted into the genome prior to the radiation of the mammalian species (**Figures 2**, **4**, **6**, **7**). Given the well-known features of retrotransposons, de-repression and transcriptional activity in germ cells, this further suggests that ancient retrotransposons may have been co-opted as *cis*-regulatory elements for establishing DNA methylation for genomic imprinting.

In mammalian genomic imprinting, one of main functions of ICRs is acquiring DNA methylation during gametogenesis [1, 2]. In the case of paternally expressed genes, including mouse *Peg3*, *Snrpn*, *Plagl1* and *Gnas*, RNA Pol II-driven transcription by alternative promoters is predicted to establish the histone modification H3K36me3 on their downstream regions, which subsequently recruits DNA methylation machinery to the associated ICRs through the interaction of H3K36me3 with the PWWP domain of DNMT3A [8–10]. A similar mechanism is also predicted to be responsible for targeting DNA methylation on ICRs during spermatogenesis, as demonstrated through truncating the low-level transcription that passes through the ICR of the mouse *H19*/*Igf2* domain [17]. However, the features of these alternative promoters or transcription have not been well understood so far. According to the results from this study, the alternative promoters of *Peg3*, *Mest* and *Plagl1* are well conserved among all the mammalian species in terms of their relative genomic position and promoter activity, triggering transcription during oogenesis. Thus, the alternative promoters of these imprinted genes may also play similar roles in the other mammals as seen in the mouse, setting up gametic DNA methylation on the associated ICRs. In that regard, it is also important to note that this predicted function has been formally demonstrated for some, not all, of these alternative promoters. Namely, the DNA methylation on the ICR of the *Mest* domain has not been tested so far, yet the two alternative promoters, U1 and U3, appear to be specific to oocytes in terms of their promoter activity (U3 in **Figure 3A** and U1 in [20]), which should be interesting to test which one is responsible for setting up gametic DNA methylation on the ICR.

These alternative promoters have another conserved feature, bi-directional promoter activity producing the two transcripts with opposite orientations: one transcribing toward the associated imprinted genes with DNA methylation setting function, whereas the other transcribing with oocyte specificity but with no known function. One likely function would be similar roles as the transcripts toward the imprinted genes, establishing H3K36me3 mark and subsequent *de novo* DNA methylation during oogenesis. According to the results from the related studies, the transcribed regions in both directions from these alternative promoters are indeed marked with H3K36me3 and also with *de novo* DNA methylation during oogenesis (**Supplementary figure S9**) [35, 36]. On the other hand, the genomic regions beyond the extent of *Ost* transcripts, especially the long intergenic regions within individual imprinted domains, are marked with another histone mark, H3K4me3, which is known to inhibit *de novo* DNA methylation [37]. As a consequence, the intergenic regions that are marked broadly with H3K4me3 have very low levels of DNA methylation [35, 37]. In that regard, it is interesting to note that the transcript lengths and promoter strengths of *Ost* are variable among individual mammals, as seen in the different lengths of *Peg3-Ost* as well as the different strengths associated with the U3 promoter of *Mest* and the U1 promoter of *Plagl1* (**Figures 4A, 6A** and **Supplementary figures S5-7**).

The variable transcript lengths and promoter strengths of *Ost* among individual species might have potential functional consequences, since the intergenic regions of imprinted domains are known to have many putative *cis*-regulatory elements [27]. One likely consequence would be that some of these *cis*-regulatory elements might have different epigenetic profiles between individual species, which might in turn contribute to some species-specific differences in the transcription and allelic expression of imprinted genes. This should be an interesting direction to pursue with epigenetic profiling of the germ cells derived from various mammalian species.

According to the results, the alternative promoters of *Peg3*, *Mest*, and *Plagl1* as well as the promoters of *Snrpn* and *Airn* might have originated from the retrotransposons that have inserted into the genome prior to the radiation of all the mammalian species. Detailed analyses indicated that these inserted repeats are the members of the ancient retrotransposon families with their evolutionary age being ∼100 mya for the L1MB7 of *Peg3* and *Mest*, ∼130 mya for the MIR of *Plagl1*, ∼150 mya for MER5 of *Snrpn*, and ∼150 mya for the L1MEh of *Airn* [29, 38]. These estimated ages are again consistent with the presence of these elements in all the mammalian species, since these insertions are likely to have occurred prior to the split of mammalian species, around 60-80 mya [39]. Then, one outstanding question would be what evolutionary forces have been driving this frequent association between genomic imprinting and retrotransposons. So far, the only known function of the alternative promoters is establishing gametic DNA methylation through the RNA Pol II-driven transcription mechanism. This may require promoter activity for RNA Pol II transcription but with strict germ cell specificity, yet the strength of this promoter should be robust and dominant to outcompete the other promoters in the adjacent genomic regions, including the main promoters of imprinted genes. On the other hand, it is well known that retrotransposons have evolved to be de-repressed and transcriptionally active in germ cells for their survival [40, 41]. Thus, the unique properties associated with germ cells might have been co-opted for genomic imprinting with retrotransposon becoming or providing a temporary strong promoter for DNA methylation setting on the promoters of imprinted genes during gametogenesis [42]. In that regard, it is relevant to bring up two individual genes, *Slc38a4* (Sodium-coupled neutral amino acid transporter 4) and *Impact* (Imprinted and ancient) which have become imprinted only in some species of the rodent lineage [43, 44]. According to the recent studies, gametic DNA methylation and subsequent imprinting of these genes is also mediated through the RNA Pol II-driven transcription mechanism involving recently inserted retrotransposons as alternative promoters for the downstream main promoters [45]. This is reminiscent of the close association of ancient retrotransposons with the alternative promoters of the other imprinted genes described in this study. Yet, these recent retrotransposons still maintain intact sequence structure compared to the heavily decayed sequence structure observed from the ancient ones, which might further allow us to identify potential *cis*-regulatory elements crucial for the imprinting of these loci [46–49]. Overall, these observations suggest that retrotransposon may have been a big part of how genomic imprinting has arisen and evolved in the mammalian genome.

## Acknowledgements

I would like to thank Drs Bambarendage ‘Pini’ Perera and Kyudong Han for their thoughtful insights regarding retrotransposon as well as feedback for the current manuscript. I would also like to thank Dr Pini Perera for her earlier contribution to identifying alternative promoters.

## Supplementary data

**Supplementary file S1. A list of RNAseq and ChIPseq data used for the identification of alternative promoters of imprinted genes.** RNAseq and ChIPseq data analyzed for the current study are presented with the original publications along with the associated SRA numbers.

**Supplementary file S2. Sequence and position information of oligonucleotides.** The oligonucleotides for RT-PCR are presented along with the relative positions within individual exons. The genomic sequences and positions of individual exons are also included.

**Supplementary figure S1. Original images of RT-PCR products separated on agarose gel electrophoresis.** All the transcripts described in the current study were analyzed with RT-PCR using a panel of cDNA prepared from the individual tissues of adult and embryo of the mouse, and the subsequent PCR products were separated on 2% agarose gel electrophoresis. The original images of these gel electrophoresis are presented along with the names of individual transcripts. The cropped images from these gel pictures have been used for the main figures (Fig 1, Fig 3, Fig 5). The PCR conditions, the annealing temperature and the cycle number for each PCR, are also presented next to the name of each transcript.

**Supplementary figure S2-S4. Promoters for mouse *Peg3*, *Mest*, and *Plagl1* in the prospermatogonia.** RNAseq data (SRR19178590/GSE236417) derived from mouse prospermatogonia was processed and the final mapped transcriptome was uploaded and visualized with IGV for manual inspection of the *Peg3* (**S2**), *Mest* (**S3**), and *Plagl1* (**S4**) locus.

**Supplementary figure S5-S7. Alternative promoters for cow *Peg3*, *Mest*, and *Plagl1* in the M2 oocytes.** RNAseq data (SRR14857179/GSE178436) derived from the M2 oocyte of cow was processed and the final mapped transcriptome was uploaded and visualized with IGV for manual inspection of the *Peg3* (**S5**), *Mest* (**S6**), and *Plagl1* (**S7**) locus.

**Supplementary figure S8. Ancient repeat elements associated with the promoters of cow *Snrpn* and *Airn.*** The genomic regions surrounding the promoters of cow *Snrpn* (**A**) and *Airn* (**B**) were analyzed with the Repeat Masker program. The transcriptional direction of both genes is indicated with an arrow. The ancient repeat elements closely associated with the promoters of these two genes are indicated with vertical arrows along with the names of repeat elements.

**Supplementary figure S9. H3K36me3 and H3K4me3 profiles of mouse *Peg3* in the full-grown oocytes.** The ChIP seq data of H3K36me3 (SRR695635/GSE112835) and H3K4me3 (SRR695641/GSE112835) from the full-grown oocytes of mouse were processed with a bioinformatic pipeline involving bowtie2, and the result of these mapping were uploaded as a sorted bam file onto IGV. The mapping result from H3K36me3 ChIPseq is shown in the upper panel and the mapping result from H3K4me3 ChIPseq in the lower panel. The transcribed region by the U1 promoter of *Peg3* is mainly marked with H3K36me3, whereas the intergenic region beyond the transcribed region by the U1 promoter is marked with H3K4me3. The DNA methylation levels show differences between the two genomic regions: the H3K36me3-marked region with DNA hypermethylation versus the H3K4me3-marked region with DNA hypomethylation.

