## Supplementary file S1 for "Ancient retrotransposon-derived promoters for mammalian genomic imprinting"

**S1 File**

**A list of RNAseq and ChIPseq data used for the identification of alternative promoters of imprinted genes**

**Mouse: M2 oocytes**

**GSE70116** (RNAseq from oocytes)

SRR2072742

SRR2072743

SRR2072744

SRR2072745

Veselovska L, Smallwood SA, Saadeh H, Stewart KR et al. Deep sequencing and de novo assembly of the mouse oocyte transcriptome define the contribution of transcription to the DNA methylation landscape. *Genome Biol*2015 Sep 25;16:209. PMID: 26408185

**GSE112622** (H3K36me3 ChIPseq from oocytes)

SRR6929776

Brind'Amour J, Kobayashi H, Richard Albert J, Shirane K et al. LTR retrotransposons transcribed in oocytes drive species-specific and heritable changes in DNA methylation. *Nat Commun* 2018 Aug 20;9(1):3331. PMID: 30127397

**GSE112835** (H3K36me3 and H3K4me4 ChIPseq from oocytes)

SRR6956535 for H3K36me3

SRR6956541 for H3K4me3

SRR6956541 for H3K27me3

Xu Q, Xiang Y, Wang Q, Wang L et al. SETD2 regulates the maternal epigenome, genomic imprinting and embryonic development. *Nat Genet* 2019 May;51(5):844-856. PMID: 31040401

**GSE129729** (H3K9me3 ChIPseq from oocytes)

SRR8889383 for H3K9me3

Sankar A, Lerdrup M, Manaf A, Johansen JV et al. KDM4A regulates the maternal-to-zygotic transition by protecting broad H3K4me3 domains from H3K9me3 invasion in oocytes. *Nat Cell Biol* 2020 Apr;22(4):380-388. PMID: 32231309

**Mouse: testis and prospermatogonia**

**GSE236417** (RNAseq from prospermatogonia)

SRR19178590

SRR19178591

Liao J, Song S, Gusscott S, Fu Z et al. Establishment of paternal methylation imprint at the H19/Igf2 imprinting control region. *Sci Adv* 2023 Sep 8;9(36):eadi2050. PMID: 37672574

**GSE236416** (H3K36me3 and H3K36me2 ChIPseq from prospermatogonia)

SRR25117953

SRR25117954

Liao J, Song S, Gusscott S, Fu Z et al. Establishment of paternal methylation imprint at the H19/Igf2 imprinting control region. *Sci Adv* 2023 Sep 8;9(36):eadi2050. PMID: 37672574

**GSE44654** (RNAseq from testis of neonates)

Li XZ, Roy CK, Dong X, Bolcun-Filas E et al. An ancient transcription factor initiates the burst of piRNA production during early meiosis in mouse testes. *Mol Cell* 2013 Apr 11;50(1):67-81. PMID: 23523368

**Mouse: the other tissues**

**GSE75957** (RNAseq from embryonic stem cells)

SRR3085918

SRR3085920

Andergassen D, Dotter CP, Wenzel D, Sigl V et al. Mapping the mouse Allelome reveals tissue-specific regulation of allelic expression. *Elife* 2017 Aug 14;6. PMID: 28806168

**Rat: M2 oocytes**

**GSE112622** (RNAseq and H3K36me3 ChIPseq from oocytes)

SRR6929782

SRR6929784

Brind'Amour J, Kobayashi H, Richard Albert J, Shirane K et al. LTR retrotransposons transcribed in oocytes drive species-specific and heritable changes in DNA methylation. *Nat Commun* 2018 Aug 20;9(1):3331. PMID: 30127397

**Rat: testis**

**GSE128824** (RNAseq from fetal testis)

SRR8783787

SRR8783788

Johnson KJ, Passage J, Lin H, Sriram S et al. Dioxin male rat reproductive toxicity mode of action and relative potency of 2,3,7,8-tetrachlorodibenzo-p-dioxin and 2,3,7,8-tetrachlorodibenzofuran characterized by fetal pituitary and testis transcriptome profiling. *Reprod Toxicol* 2020 Apr;93:146-162. PMID: 32109520

**Human: M2 oocytes**

**GSE157834** (RNAseq from oocytes)

SRR12642761

SRR12642763

SRR12642767

Asami M, Lam BYH, Ma MK, Rainbow K et al. Human embryonic genome activation initiates at the one-cell stage. *Cell Stem Cell* 2022 Feb 3;29(2):209-216.e4. PMID: 34936886

**Human: testis**

**GSE144085** (RNAseq from testis)

Tan K, Song HW, Thompson M, Munyoki S et al. Transcriptome profiling reveals signaling conditions dictating human spermatogonia fate in vitro. *Proc Natl Acad Sci U S A* 2020 Jul 28;117(30):17832-17841. PMID: 32661178

**Crab-eating macaque: M2 oocytes**

**GSE233232** (RNAseq from oocytes)

SRR24706082

RNA-seq for the comparison of gene expression between monkey GV oocytes, in vitro matured oocytes and in vivo matured oocytes.

**Crab-eating macaque: testis**

**GSE95736**

SRR5314740

Clark AT, Gkountela S, Chen D, Liu W et al. Primate Primordial Germ Cells Acquire Transplantation Potential by Carnegie Stage 23. *Stem Cell Reports*2017 Jul 11;9(1):329-341. PMID: 28579394

**GSE69241**

SRR2040595

Ruiz-Orera J, Hernandez-Rodriguez J, Chiva C, Sabidó E et al. Origins of De Novo Genes in Human and Chimpanzee. *PLoS Genet* 2015 Dec;11(12):e1005721. PMID: 26720152

**Cow: M2 oocytes**

**GSE178436**

SRR14857179

Cuthbert JM, Russell SJ, Polejaeva IA, Meng Q et al. Comparing mRNA and sncRNA profiles during the maternal-to-embryonic transition in bovine IVF and scNT embryos†. *Biol Reprod* 2021 Dec 20;105(6):1401-1415. PMID: 34514499

**Cow: testis, hypothalalmus, adrenal gland**

**GSE176219**

SRR14740596

SRR14740612

SRR14740620

Rabaglino MB, Bojsen-Møller Secher J, Sirard MA, Hyttel P et al. Epigenomic and transcriptomic analyses reveal early activation of the HPG axis in in vitro-produced male dairy calves. *FASEB J* 2021 Oct;35(10):e21882. PMID: 34460963

**Pig: M2 oocytes**

**GSE217367**

SRR22199132

Murin M, Nemcova L, Bartkova A, Gad A et al. Porcine oocytes matured in a chemically defined medium are transcriptionally active. *Theriogenology* 2023 Jun;203:89-98. PMID: 37001226

**Pig: testis**

**GSE233410**

SRR24735911

Wang P, Liu Z, Zhang X, Huo H et al. Integrated analysis of lncRNA, miRNA and mRNA expression profiles reveals regulatory pathways associated with pig testis function. *Genomics* 2024 Mar;116(2):110819. PMID: 38432498
