## Supplementary file S2 for "Ancient retrotransposon-derived promoters for mammalian genomic imprinting"

**S2 File**

**Sequence and position information of oligonucleotides.**

The oligonucleotides are indicated with underlines within the exons of Oocyte-Specific Transcripts (OSTs).

***Peg3***

>Exon-U1 (chr7:6,751,059-6,751,235 in mm10)

GCAGGGAAGCCTCCATCCGTTTGTCTTTAGGCTTTCCATTCCCAGCATCTCATAGAAGTGTAATCTTCGCTATTTACCATCCCTTCTTTTAACTAATCAAGGTTGCAGTTTTATCAGGAGTTTGGTATCCCTCTTAAATGGTTTGAGAAGCTCAGAACTGCACTGCAGGGGATTCAG

>Exon-U0 (chr7:6,746,702-6746,839 in mm10)

ATACAATGATACAACATGGCCAGCCTTAACCACAGATATAATGATGCAACATGACCAGCAGTTCCTCCCACTCCAATTGAAGCCCTGAGCTCCTGCACACTCCAGCTTCACTGCCATGATGGACCCTACCCTGGAACT

>Exon1

AGACGCTGGGGAGTCAGGAGTCGCGGGAGGACGAGCATCGGAGGAGAAGCGGAGAGATGTCCACCCTGGGCTGGTGGCGCCGCCGGGCGCCCGGTTCAGTGTGGGTGCACTAGACTGCCGACCCTGGTCGGGGTGTGTGCGTAGAGTGCTGTGCTCCGGGAG

>Exon2

CCCTACCTTCTTGATCTTCTATCCTTTTTGGAGACAACTGGCAAGAGGAAGACTAGGTCCTCCAACAGCTAGGTGGCTGGTCCAG

>Exon3

GCAGGCCTTCCCAGCAAGGGGAGATCAGTTGATCATCCCTGAAACGCTCAAGCCCTTGGGTGTGAGCAAAACAGACAACTGTGAAAAACTCACCACTCCGTTGGAGAGTTTCAAGATGTACCATCACGAAG

Primers for RT-PCR

>Peg3-RT-U1-F

AGGGAAGCCTCCATCCGTTTGTC

>Peg3-RT-1a

GGTTCAGTGTGGGTGCACTAGACT

>Peg3-RT-1b

GCTCACACCCAAGGGCTTGAGCG

***Peg3_Ost***

>Exon1 (chr7:6,751,416-6,751,595 in mm10)

GTATGCTTGGTGCAGTTTTGACTTCCGTGAATCAAATCTCAATAAAGCTTTAAAGGTCAGTTACCTTGGCAAAGAATCCTCAGACCCCTCGCTAAGCCTCTTAGGGTCTTCTCCTCCTCTGGTCGATGCTAATTAGGAACGCCAAGTTCTGAGAGAGAACACAGCAGTTGTAGCGCAAAG

>Exon2 (chr7:6,751,570-6,751,598 in mm10)

GTAGCCACATCTGCTGTGTGTCCATGATTGCATGCCATGTCCAGAAGACGGTGTTTCATGTCGCACCTTCCCATCCTGCCATCAGCCCTACGTCCTACACAGTAACTCAGCTGTTCCCTGTACAGGTCTCCCCCATGCCCCTTCATCTCCATGTCTCCATGGCTGTACCTGTGTTCCTGGTCACCCTGTGTCTCCCTCACTCCCCGACTTGAGTGGCCTCTGATGAACTCAGCACTTGGCAGCCACAGGGATCTTCTCAAGAGGAAAATTGGAGGGCATCACCTCCTTGCTTGAAACTGGCCAGTGCCCATTAGACTGAAGACCACGCCTCTCAGCATTCGCCTGACTGTCCAGTAGGCCATCAACCTCCTCTGTTATTGGTTTCCCCAACATCAGTTTATTTCATCTTCCTAAGCCCCAGTGGAATGGATATTCTAAGAGTTCCTGCCTCAGTCAAAGGGCTTGCTTCAGAATCAGGAGAACATGAGTTCAATGCCCAGAACTATGTTAATGACAAACAAGCAAACAGACCAACAAAATCTGGGGGTTGGGGGGTGGGGGGAGGAGAGGCGGGTGCTGTGCACTTGTAACCCCAGTGCTGCCAAAATAGAGCAGTAGGGATCTTTGGCCAGGTAGCCTAACCTACTTGTTGAGTTCTAGGCCACTGAGAGATTCTGTGTTTTTTTCTTTTTTCTTTTTTTCTTTTTTTCCTTTTAAGGTGAATAGTGACTGAGAATAGCATCTGAGTTTGCCCTTTGGCTTCCATGAACATGTGCACCCAAGGGCACCCTCAGCTCTACCACCCCTATATACAAACACACAGAGATTGTATCTTGAAGGAGATCGTCTGTACAGAGTCTTAGCAGAGGAATCTTTAAGGGCTGTTTTGTAAGCATAGTTCAAAGTTCCCGAATTCCAGACACTATGCTTGGGCAACAGTCATTTGGAAAAACAAGTTCTTGGTTGTCTGTATCCAGCAGTTCCACTCTGGACATAGACCCAAAAGAAATGAAAACAGGGCCTTGAAGATTTTGTTATGCTTGCAGTCATACAAGCATTATGTATGCCAATCTAGGGGCAGAAAAGAGCCACGTGTTTATCTATACATGAACAAATAAAATGTGGCATGCCCATTGAGTGGGATACTATTCAGTCACAAAAAGGGAGGAAATTCTGACACATGCTCTGTGTGTAAAAGTCCAGGACATTAAGCTAAGTAGGAAAGGAGCAGACACCAAAATGACAAATGCTATGTGATTCCAGTTATTTCTGGCAGTCAAATCCTTGGATGTATAAATAAGGATGGTGGTTCCAGGTGTTCTTCCAGCAGCTGCCACCTCCTCTTCTTTCTCCTCCTCCTCCTCCTCCTCCTCCTCCTCGTTTTCTGCTTCTTTTCTTTTAACTTATTTATTTATTATTGTATGTGTGTGGCTTCATGCATACACATGTGTCTTGGCACATGTAATAGAAGTGAGAGGACAACTTTCAAGAGTTGCTTCTTTCCTTCTACCTTGTAGGGATTGAACTCAGGTCATTAGACTTGACTAGCAAGCTCCTCTACCATCTGAGCCATCTTGCTGACCCTGCAGCCATCCTAATTGGTTGGTGATGCTGCATCATTGTCTCAGATTGTAATTTTCTAGTGATTAATGGCATCAGATGACTTTTTATGCGACTGTTGGCCATTTTATATCTTTGGAGAGACAGTAATTTAATGTCTTTTTCATTTTTAAGAGATAAAACGGAGTTGTTTTTTTTAAGATTTATTTTATTGTGAGTGTTCTGGTTGCATCTGTGTCTGTGTACTACATGTGTGCCTGGTGCCCATGGAGGCCAGAAGAGGGTGATGGATGCCCTGGAACTGGGGTTACTGGTGGTTGTGAGCTACCTTGTGGGTGCTGGGGACTGATCCACAGTCCTTTGGAAGAGAGAAGCCAGTACCTTCACTGCCAAACCATCTCTTCTTCCACATTACTTTGCTCCTTCAAATTGTTGAGGTCTTGGAGTCCCCTTATATATTTAAACTGCAGTGCTTTATGAGGCACTGCCCCTCATTTTATAGATTGCTGTTTTAAAATGTTATGAGTGTGTACAGCATATGCTGGAGCACTGTGGTACACGTGTAGAGGGCAGAGAACAACTTTTGAAATGTCTTTCTTTCCTTCCATCGTGTGAGCAGAGCTCAGGTTGGGAAAGCACCTACCAGTGATCTCACTGCCTCCCTCCCACCCCCCACCCCCCAGCCCCAAACTGCCTTTGTAGTCTGTCAACCATTGTATCCTATGGCCATAGAAGGTTTTTTTGGTTTTTGTTTTGTTTTGTTTTTTTAAGATTGATGTAATTCTATTTGTCTGTTTTCACATTCTTCCCTGTGTTTTCTTCGGCATGTGTGTGAGCGTAGTATCACGTGACTCCATTCATTTTTTTTTTTTTAAATTTTATTTTCTTTGCAGTACTGAGGATTGAACCCAGGGCCTGTGCAGGCTAGGTGAGTGCTCTACCAACCGTGCCACAGCATCAGATCCCTATTCACTTTTTAAACTTCCTTGCCTTGATTGCCCATATCTTGCTCCCTGCTCTATCACTTCCAGGTAAAGAGCCTGACTTTTCTATGGTGATGTGTGGCCATTGAGATAGGGTCTCACCAAGTCACCCATGCTGGCCTGGAACTCACAGTCCTCCTACCTCAGTCCTCAGAGCGCTGGACACAGGCCTGCATGACCAACCACTGCCAGGTAGCAGGGGCCTGATTCTGTTTATTCCCTGATCCCTGAATATAAGCTGTGACTGAGTACTCCATTGCTAGTGTTTGACAAGATTTGACTTAGAGTTTACACCTGACCATCCCTGGGCAGTGTGCCTCAAGATTCTTCCTTTGTACATTATGAAAGGAGGGATGAGGGTTTCAGGAGAGGAGCCACCTGCCATTCCATAATATGACACTCAAGTGATGAGTTGAAACTGTTGATGTCTTCAGAGAGTTTGCAATGGAGCAGGGACTAGGGAGTTTCTGTGTTCTTTCTTCATGAGAAGGGACTCTCAGAAGAGGCATGAACCTGAGGCTTTTAAGAATGCTGAGGGCCATCAGAATGAGGTGCAGGAGATCTTGGCTTTATTTCTACCCTTTCTGCTGCAGCAGGAGGAAAGTTTTGAGAGTCACTTCTCCAGAAGAGAATGGGAGCCGAATCCACACCCTCAGCTACCAATATTATCTTCATGAATATGAAAATATAACCAGGGTCCACACCTTCCCTCTGTAGGATATTCTCAAACATTCTAGGTTTTATCAGACTTTCATGGAGTTCCAGAAACCTCTGGGAGGATCCGTGGACTTCTGTGCAAAATCTTATATTAGCAAGAGCCTACACCCCCACATCTGACATATCTGATCACCTTCCTAATCCCACATGGTGGCATCTCTCCTTCCAACCCCCACCCTGTGTTCAGCAAGATTGCTGGTTCAGAGTGGAGAAAATCACAATGTTTCCATGGGGGACCTTTCCTATCCATGGACCTGATCCCGACTCTTGCTTATAGATCTCCAACTCACCCTACACTGTGTTTGGAGTTACAGTGTATCTTCATCCCTCGAGCTCCATAGGAAAACCCAGGGCTGGTGAGATGGCTCAGCTGGGTAAGAGCAGGGCCACTTGAATGCCTAGGACTGGGCATTCTTTCCATCATGGAAAGGAGAGGGCTGACTCCTGCAAGTTGTCTTTTGATCTCCACAAGTGCCTATCCCCCTTCATGCAAAATAAATAAACTAATGTAATTGAAGAAGAAAAAAAATCCACTGTCA

Primers for RT-PCR

>Peg3-Ost-F1

ACACAGCAGTTGTAGCGCAAAG

>Peg3-Ost-R1

GTTCATCAGAGGCCACTCAAGT

***Mest***

>Exon-U3 (chr6:30,711,062-30,711,217 in mm10)

ACCATGGCCTCTGGAGGCCAGACAGTGGGATTGGCGCGTTGCTTGCTATCTGGTCCCTCAGAAGTGCTGGGTAAAAGACTCCCCGTGCTGCTTCAAGGCCTTGGGTAAGCCATGGCGATTGGCCAACGTGGGGACAGAGGCCCGTTGGTTGTTTAAG

>Exon-U2 (chr6:30,723,546-30,723,646 in mm10)

CATACTTTTCAGGCTACCCAGGAGCGCGGCTGCTCCCAGGGCTGCATGCGCAGCCTCCAAGCCTTCATGCATGGGCATTGGCTCTCCCAACCCAGCCACAG

>Exon-U1 (chr6:30,733,506-30,733,613 in mm10)

GGGGTAGAGAGAAAAAGTGTGGAAGGCTGCGGTCTAGTCTTCCTTCTTGGGCAGCTGGGAAGAGAAAGCCAGCTTGTTTGGAAGTCGCTGTTCCTTAGAGGGCCTGTG

>Exon1

CCAGCACATCCCGGTGCTTCTTCTCAGGCGCAGCAGCTTTCCTCTGCGGCAGCCGCACCTCGCCAAACGGCGTAGTGCTGCAGGCTCGCCCGAGTTGCTGCTTGCTGCCTCTGCTGCCGCTGCCGCGGGCCGCCCTGCGCGGACCGTAGGCTGCGCAGACGCCACCTCCGATCCTGTATCGCTGCGGGCGCCTCGGCGCGCCCTGTGATCCGCAATCCTGCGGCGGGCGGCATGGGATAATGCGGCCATGGTGCGCCGAGATCGCTTGCGCAG

>Exon2

GATGAGAGAGTGGTGGGTCCAAGTAGGGCTCCTGGCTGTGCCCTTGCTGGCTGCGTACCTGCACATCCCGCCCCCTCAGCTCTCCCCTGCTCTGCACTCATGGAAGACTTCTGGCAAGTTTTTCACCTACAAAGGCCTACGCATCTTCTACCAAG

Primers for RT-PCR

>Mest-RT-U1-F1

GGGGTAGAGAGAAAAAGTGTGGA

>Mest-RT-U2-F1

CATACTTTTCAGGCTACCCAGGA

>Mest-RT-E1-F1

GCCCTGTGATCCGCAATCCT

>Mest-RT-E2-R1

CTTGCCAGAAGTCTTCCATGA

***Mest_Ost*** (***Cep41_U1)***

>U1 of Cep41 (chr6:30,710,072-30,710,613 in mm10)

CTTCTCAGAGTGAGGTTGGCTATTTGAACTAGGGACCATAAGCTAGGAACATTCTTTATTTTATTAGTTCTTAAATTTTTAGTTTTACTTAGTTGTTGTTGTTGCTAGTGGTGGTGTGTGTACCATAGCGTTCCTGTGACGATTGGAGGACAACTTCCAGGAACCTGTTCTCTCCTTCTGTCGCCATGTGGGTCCCAGGGATTGAACTCAGGCCTTCAGGCCTCACAGTGGGCGCCTTTTATGCCTTTGTCCCCTTTGCCCAGTGAGCTATCTTGATGGCCCAGGAACATTCTTTTAACCTGCCCATATTGCTTACTTCTGTTTTCTCTCTCAGTTTAGTGTTTCGAAAATAAAATCTTTAAAAGCAAAAGCCCGCCTGGATTTCTGTGGGGCTGAGAAGAACACGCGTCTTGTCGTGGTTGTAGCTTTGCTGCTGGTTTTGAGAGGACCAGTCCTTGGTATGGAGGGGCAATTGTTAGAGGGCCATGTGCCCTTGTCCCCTGGAACAACCCAGTTTATTTTTTGTTAACACAGTGTAACCA

>Exon2 of Cep41 (chr6:30,680,131-30,680,194 in mm10)

CAGTATCCAGTCTTGACTTGACATGTTGATACCTTGGGTTCTGTGGTATCCGTCTGGTCAGATA

Primers for RT-PCR

>Mest-RT-U3-F1

CAGTGGGATTGGCGCGTTGCT

>Mest-Ost-F1 or Cep41-U1-F1

TCCTCCAATCGTCACAGGAAC

>Cep41-E2-R1

TCCAGTCTTGACTTGACATG

***Plagl1***

>Exon-U1 (chr10:13,060,505-13,060,859 in mm10)

CAGAAAACAGACCACCTGGCGAGTGTGCCAGGCATTTTAGCAAGGCTTCTCACAGGCTTAATTCTTGCAACCAGTTTAAATGAAGACTAGGGCAGGGTGTTGGTGCCTGAAGCTAATGACTATTGAGGGCTGGGCTTCCTTATTCGGAAGCCTTGGACAGCCCAGAACCTTAAATTTGTCTGGAAGATTTTCTTCCCAGCTTTTAAGCTCCAGATCCCAGAGCACATGGATCCGCTCATGGGCAGGTATCACGACCTCCCCACTTCCGCAGAAAGCGGTAAAGTTAAGCGGCCAAAGGTTGTACACAAATAAGGACCATGGAAGTATCTGCACCCAGATGGCCTAACCCCTAAAG

>Exon1

GGACCGCCCCGAGCCTTGATTTAGCCGGGGCTGGGGCGTTCTCCAACCTCACTCGCCTGGCAGGCGGGAGAACGCTCGGGGAGTTGCGGCCGCGGGCACCGGGCTCGCGGCTATCGGGACTGGAGAGCAAGCGGGCATCTCCTGGGCGCCGTCATGGCTGCTTAGGCTGCGCCTGCCTGCGGATCGCGGATCCGGGATCGGAGATCTGACGGCGACGCCTGAGTCCGGCTAGGGTAG

>Exon1.a

GTTGTTCCTGCTTGATTGCTTCAGCGTGCCATCGGCTTC

>Exon2

GTATTTGCATAGGAGTCAGAGGAGTTAATCTT

>Exon3

CTCTTCTCACAGGTTTGAGTCTTCAGACTTCTACAGAACTCCATAATATCTGCCTCACAGCTGGCTTTCCTGCTCTCACAGAAG

Primers for RT-PCR

>Plagl1-RT-3

AGCGGCCAAAGGTTGTACACA

>Plagl1-RT-4

CCTCTGACTCCTATGCAAATAC

>Plagl1-RT-2

GGTCTGGAGGTGGTTCTTCA

>Plagl1-RT-1

TTCGTCACCCTGGAGAAGTT

***Plagl1_OST***

>Exon1 (chr10:13,060,172-13,060,423 in mm10)

CAGACAGTGCTTTTCTTTGTTGTTGAAGGCTGGGGAAATTTTGCAGTGAGAGTTCAGAAACAGGTATCTCAGAGGTCCTTTGTTCTTTCTGGGGGAATTTCTTCTTCTTTCCTTTTCTTGCCCTTGCCCTCAACAAAGTCTTTTTTTTTTTTTTTTTAACATATTGCTCTGGGTGAGCTGGGACCCAGGTAAAACCAGGTACTTGTAATGTGGCAACTGAGTTCTGCCAGACCCCGGAGGAAGCTGCGTGGG

>Exon2 (chr10:13,043,314-13,044,586 in mm10)

TCTGCCCTAGTTTTCAATGATCCTTCCTGTCTGCTGCTTAATTTTCCTCATTGTTATTCCAACACATTGAAGCGTATCCTTGAGTTTTGATTTATCTCAAACATTCTAATGTAGGCAGTTAGCTTCCCTCTTAGGACTGCATCGTGACCTCAAGAGTTTGTTCATTCTTTGTTAAATTCTAGTGTATTCTTCAGTGTTGGGGTAAAATGCTCTGTCGATAACTGCTAGGTCCATTTGATTCATGATGTCTCTTCATGTACCGCCCCCCCCCCCCAAGCCTGAAATCGGTCTGCCACTCCCACTGCTGCTACCTGCCGCCTAGCTGTTCCGGACTGCAACTGGACTCCAAGGATGATTTGGTGGGAATGGGCTCCTTCCCCCTTCTTCATAACCCGGTGCTCTGAACAGTAAAATGGAGCTTTGATCAGAACCTTGTTGTCTTAGCTCCATTCTTTCTCTCATCTGTCTAGTTCCCTTTCTTTCAGCTTGATTCTGCCGCTTCAGCCATATCCCGTTCCTCCCAAGCCGCTGGACGACATGCAACATCTTCACTTCAGTGTTTCTCTAGTTACTTTTTAACATGTGGATACTTTGTCTATTGGTGAGGGTGGGGTACTGAAGCCCCCCACTATCACCATGCTGGGGGCTTGGATCTGGTAGCATTTGCTCTATGAAAGTGGGTACATGTTTAGAATTGCAATGTCTTCTTGGTGAACTGTTCCTTTCATCACTATGAAGTGACTCTTATCTTTTCTTGTGAGTTTTGATTCTGTCTATTTTTAAAGCAGCTATTAGACTAGCAATGTCTGCTTGTTTCCTGGTTGTATTTGCTTGGGATAACTTTCTATCCTTTTAATCTGAGATAGTATCTACTGTTGATGGCAAGATGTGTTTCTTGGAGGCAGTAAGGAGAAGGCTCCTGTTTCCTATACCCATCTGTGAGTCTGTGACTATTTATTGGAGGCATTGAGAATATTGATATTCGGCGAGTATTCAGTTGTGTCTCGATTCATGTCGTTTTGTAGATTTTGTGGCATTTCTGTAGACATCTTTTGCTTTCTTTCAGTCCTTGGAATACATCTTCCTAGGTGTTTCTGGCCTTCAAAGTCTCTGTCAAATAATTGGTTGTTGTTCTAATGGGCCAGACTTTATAAGGGACCCAACCTTTGTAGCTTTTCAATAACCTTTTAGTTTGTGTTTTTGATGTTTTGACCATATGTTGTAGGGGAATTTCTTTTCTCGTCCTATCCATTTTTCTGTAGGCCTTGTATAC

Primers for RT-PCR

> Plagl1-Ost-F1

CCTGTTTCTGAACTCTCACTG

> Plagl1-Ost-R1

GGGCCAGACTTTATAAGGGAC
