## Supplementary figures and images for "Ancient retrotransposon-derived promoters for mammalian genomic imprinting"

### Supplementary figure S1

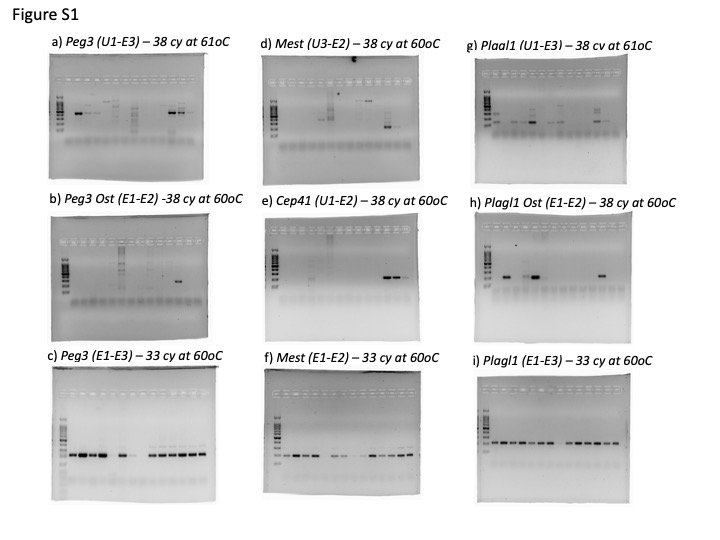

### Supplementary figure S2

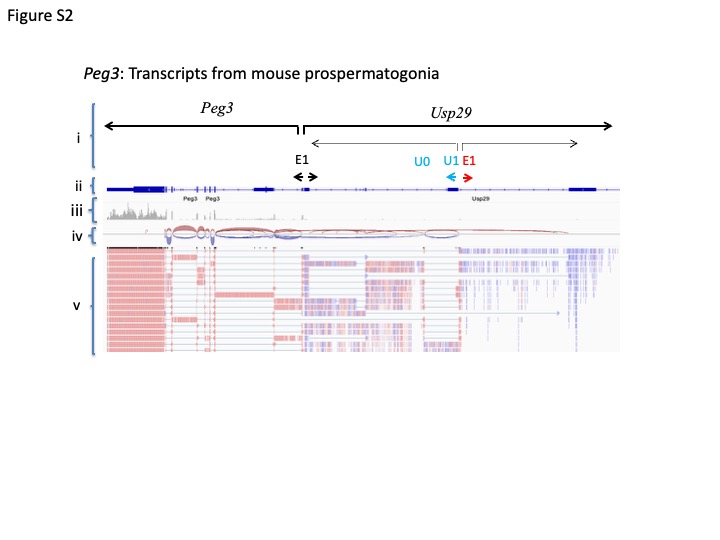

### Supplementary figure S3

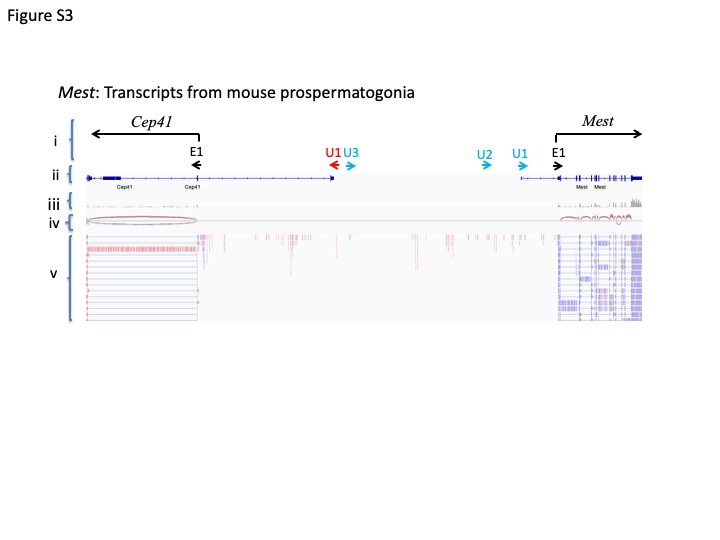

### Supplementary figure S4

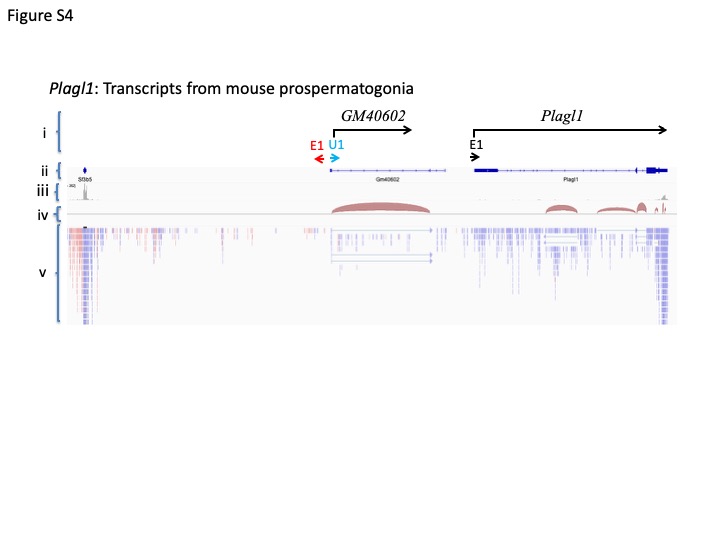

### Supplementary figure S5

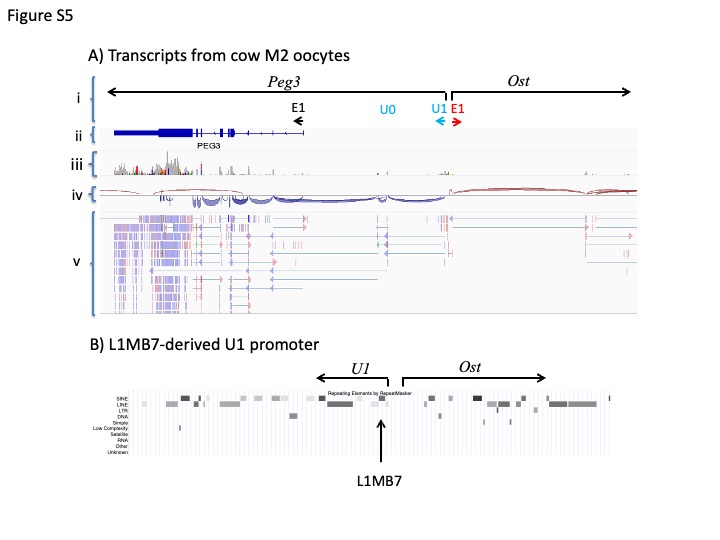

### Supplementary figure S6

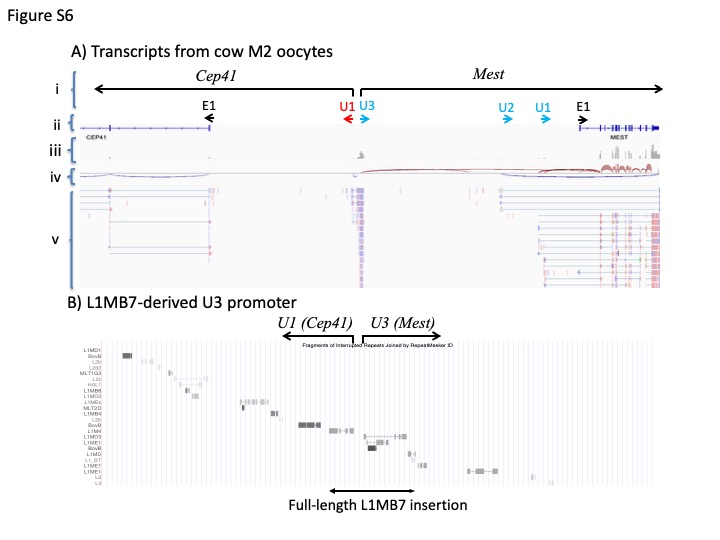

### Supplementary figure S7

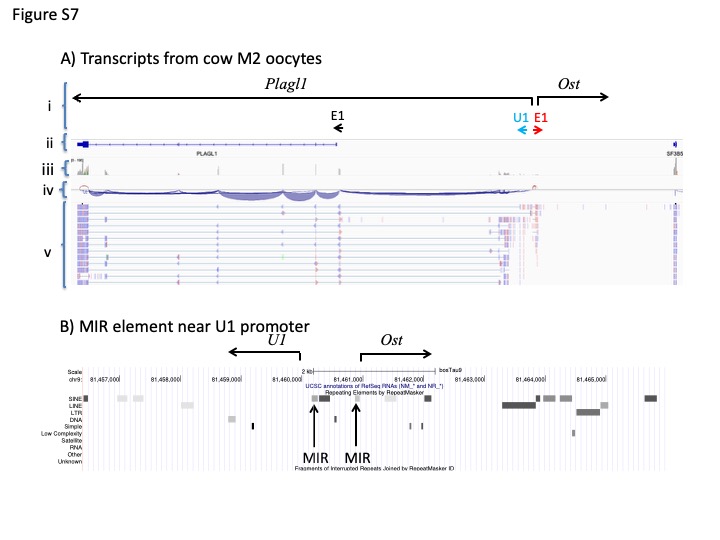

### Supplementary figure S8

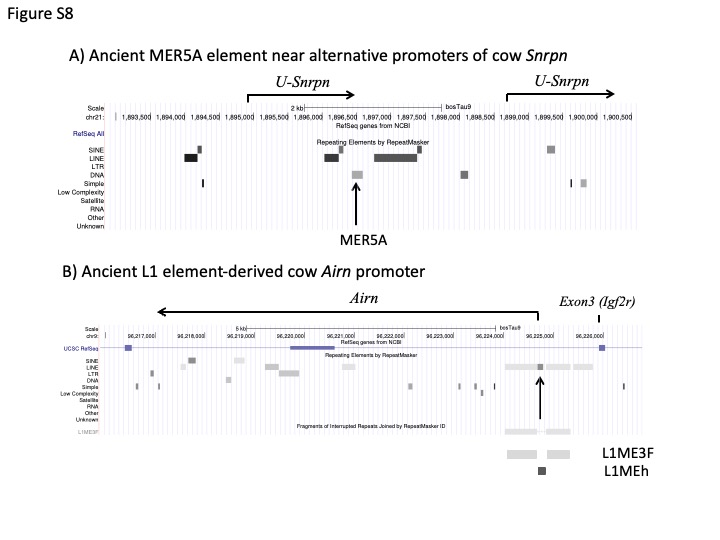

### Supplementary figure S9

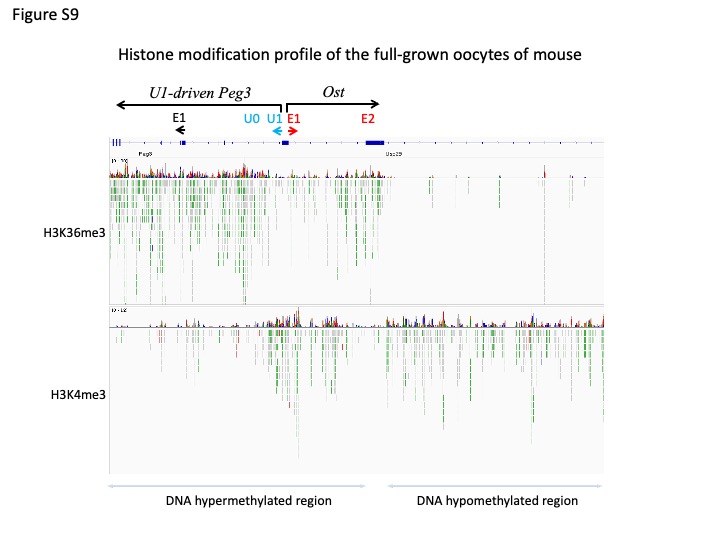
